# Infection-experienced HSPCs protect against infections by generating neutrophils with enhanced bactericidal activity

**DOI:** 10.1101/2022.11.03.515114

**Authors:** Hannah Darroch, Pramuk Keerthisinghe, Yih Jian Sung, Anneke Prankerd-Gough, Philip S. Crosier, Jonathan W. Astin, Christopher J. Hall

## Abstract

Hematopoietic stem and progenitor cells (HSPCs) respond to infection by proliferating and generating in-demand neutrophils through a process called emergency granulopoiesis (EG). Recently, infection-induced changes in HPSCs have also been shown to underpin the longevity of trained immunity, where they generate innate immune cells with enhanced responses to subsequent microbial threats. Using larval zebrafish to live image neutrophils and HSPCs we show that infection-experienced HSPCs generate neutrophils with enhanced bactericidal functions. Transcriptomic analysis of EG neutrophils uncovered a previously unknown function for mitochondrial reactive oxygen species in elevating neutrophil bactericidal activity. We also reveal that driving expression of zebrafish C/EBPβ within infection-naïve HSPCs is sufficient to generate neutrophils with similarly enhanced bactericidal capacity. Our work suggests that this demand-adapted source of neutrophils contributes to trained immunity by providing enhanced protection towards subsequent infections. Manipulating demand-driven granulopoiesis may provide a therapeutic strategy to boost neutrophil function and treat infectious disease.

## INTRODUCTION

We now know that innate immunity has the ability to ‘remember’ previous microbial encounters and generate stronger responses to subsequent infections with the same or unrelated pathogen (Netea et al., 2015; Netea et al., 2011; Netea and van der Meer, 2017). This memory function, termed trained immunity, is underpinned by epigenetic and metabolic changes within innate immune cells that support elevated antimicrobial responses. The longevity of trained immunity is the result of similar infection-induced changes within hematopoietic stem and progenitor cells (HSPCs) that provides a longterm supply of trained innate immune cells (Christ et al., 2018; Cirovic et al., 2020; Kaufmann et al., 2018; Mitroulis et al., 2018). The adaptive ability of HSPCs to sense bacterial threats and generate in-demand innate immune cells (termed demand-adapted hematopoiesis) is now recognized as an integral component of trained immunity. Further connecting demand-adapted hematopoiesis and trained immunity is the transcription factor C/EBPβ, a central regulator of emergency granulopoiesis (EG; the production of in-demand neutrophils following infection) that has recently been shown to also control the epigenetic reprogramming of HSPCs during trained immunity (Akagi et al., 2008; de Laval et al., 2020; Hirai et al., 2006; Sato et al., 2020).

To date, studies describing trained immunity have focused almost exclusively on monocytes/macrophages. Being the most abundant circulating leukocyte, neutrophils provide the essential first line of defense against microbial challenges. Fundamental neutrophil effector functions include phagocytosis and the subsequent destruction of intracellular bacteria through reactive oxygen species (ROS) production (Kolaczkowska and Kubes, 2013). Only recently have neutrophils been shown to contribute to trained immunity. Neutrophils from humans vaccinated with BCG (a driver of trained immunity) have been shown to respond more efficiently to microbial challenge (Moorlag et al., 2020). Furthermore, treating mice with β-glucan (another agonist of trained immunity) skews hematopoiesis towards the production of neutrophils with enhanced anti-tumor activity (Kalafati et al., 2020). Although these studies have revealed neutrophils as mediators of trained immunity, mechanisms that instruct their trained phenotypes remain poorly defined. Furthermore, whether EG contributes to trained immunity through generating neutrophils with heightened antibacterial responses is unknown.

Equipped with a highly conserved innate immune system and supported by neutrophil- and macrophage-marking transgenic reporter lines, transparent larval zebrafish provide a powerful system to directly observe the host response to microbial infection, at the single cell level (Linnerz and Hall, 2020; Torraca and Mostowy, 2018). Larval zebrafish can also model EG, where large numbers of neutrophils are produced within larval hematopoietic sites, the aorta-gonad-mesonephros (AGM) and caudal hematopoietic tissue (CHT), following infection (Hall et al., 2012; Hou et al., 2016; Keightley et al., 2017; Liongue et al., 2009; Willis et al., 2018). In addition, their exclusively-innate immune system when at larval stages has been exploited to model trained immunity (Darroch et al., 2022; Willis et al., 2018). We, and others, have shown that larval zebrafish HSPCs demonstrate a conserved response towards a sub-lethal bacterial challenge by expressing the zebrafish ortholog of *C/EBPβ (cebpb),* expanding in number and enhancing their contribution towards the neutrophil lineage (Hall et al., 2012; Willis et al., 2018). Recently we showed that this EG infection model can drive an overlapping trained immune response (Darroch et al., 2022). A feature of our EG model was a cohort of infected larvae that became almost completely neutropenic prior to *de novo* neutrophil production (Hall et al., 2012). We speculated that this would facilitate the study of EG-generated neutrophils in isolation from those generated under steady-state (SS) conditions, for the first time within a live animal model.

Here we provide evidence that infection-experienced HSPCs drive an EG response that populates the host with neutrophils equipped with enhanced bactericidal functions. Our work suggests that this demand-adapted source of neutrophils likely contributes to trained immunity through providing enhanced heterologous protection towards subsequent bacterial challenges.

## RESULTS

### Larvae populated with EG neutrophils demonstrate enhanced survival to subsequent infections

To generate larvae populated with EG neutrophils, 2 day post fertilization (dpf) neutrophil-marking *Tg(lyz:DsRED2)* larvae (Hall et al., 2007) were injected with a sub-lethal dose (600 CFU) of GFP-expressing *Salmonella enterica* (hereafter referred to as *Sal-*GFP) into the hindbrain ventricle, as previously described (Hall et al., 2012). Infected larvae were then manually inspected for neutropenia at 1 day post injection (dpi) by fluorescence microscopy (Figure 1B). Here, neutropenic larvae are defined as those that possess no more than 15-20 remaining neutrophils. Larvae with high neutrophil abundance, or that retained residual *Sal*-GFP (as assessed by fluorescence microscopy), were discarded. Larvae that underwent EG were selected at 2 dpi (4 dpf) by identifying those with *de novo* neutrophil production within the AGM and CHT (Figure 1D). On average, 27.9 ± 1.5% (mean ± SD, n=~100 larvae in biological triplicate) of infected larvae became neutropenic, and of those larvae 74.5 ± 3.9% demonstrated strong EG. To generate larvae populated with SS neutrophils, 2 dpf *Tg(lyz:DsRED2)* larvae were injected with PBS (Figure 1A and C). Throughout the remainder of this study, larvae populated with SS and EG neutrophils are referred to as SS and EG larvae, respectively.

**Figure 1.**
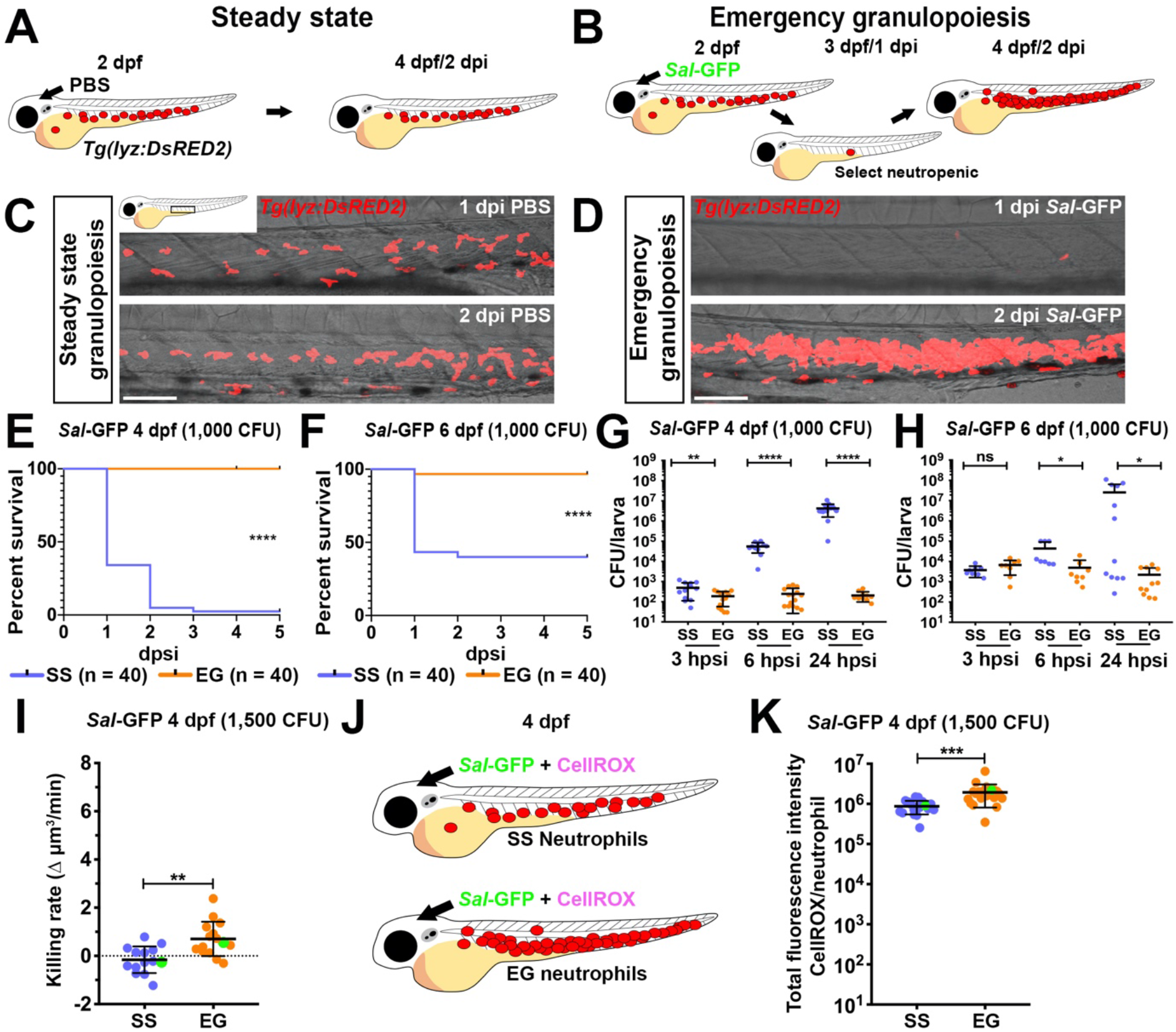
EG larvae have elevated survival to subsequent bacterial challenge and possess neutrophils with enhanced bactericidal activity. (A and B) Schematic illustrating strategy to generate SS (A) and EG (B) larvae. (C and D) Live imaging of the AGM/CHT regions within *Tg(lyz:DsRED2*) larvae at 1 and 2 dpi with PBS (C) or *Sal*-GFP (D). (E and F) Kaplan-Meier graphs showing survival of SS and EG larvae over 5 days post secondary injection (dpsi) with *Sal*-GFP at 4 dpf (E) and 6 dpf (F). (G and H) Bacterial burdens within individual SS or EG larvae at 3, 6 and 24 hours post secondary injection (hpsi) with *Sal*-GFP at 4 dpf (G) and 6 dpf (H). (I) Bacterial killing rates of SS and EG neutrophils following *Sal*-GFP infection. Green data points highlight killing rates of neutrophils as shown in Figure S1J. (J) Schematic illustrating injection of SS and EG larvae with *Sal*-GFP and CellROX. (K) Quantification of ROS production within *Sal*-GFP-laden SS and EG neutrophils, as detected by CellROX fluorescence. Green data points highlight ROS production of neutrophils as shown in Figure S1M. Error bars, mean ± SD; ns, not significant, * p<0.05, **p<0.01, ***p<0.001, ****p<0.0001; Gehan-Breslow-Wilcoxon test (E and F), unpaired Student’s t-test (G, H, I and K). CFU, colony-forming units. Scale bars 50 μm.

To initially investigate differences between SS and EG neutrophils we infected SS and EG larvae and monitored their survival. SS or EG larvae were infected with a lethal dose (1,000 CFU) of *Sal*-GFP into the hindbrain ventricle at 4 dpf and monitored for 5 days post secondary injection (dpsi). This analysis revealed that EG larvae had a significant survival advantage that was associated with enhanced bacterial clearance (Figure 1E and G). Furthermore, the enhanced antibacterial response of EG larvae was maintained when extending the interval between primary and secondary infections to 4 days (Figure 1F and H). To better assess the contribution of EG neutrophils to the enhanced survival demonstrated by EG larvae, we injected liposomal-clodronate (Hall et al., 2018) into SS and EG larvae (at the neutropenic stage) at 3.5 dpf, prior to *Sal*-GFP infection at 4 dpf (Figure S1A). This treatment resulted in ~80% macrophage depletion at 4 dpf (Figure S1B) while leaving neutrophil numbers (Figure S1C) and the EG response (Figure S1D) unaffected. As expected, macrophage-depleted larvae were more sensitive to *Sal*-GFP infection (Figure S1E). However, macrophage-depleted EG larvae still maintained a survival advantage, when compared to macrophage-depleted SS-larvae (Figure S1E), a phenotype supported by significantly enhanced bacterial clearance at 9 hours post secondary injection (hpsi) (Figure S1F and G). Of note, despite EG larvae possessing more neutrophils than SS larvae (~2.5 fold increase [Figure S1D]), similar numbers were recruited to the infection site (Figure S1H and I).

### EG neutrophils have enhanced bactericidal capacity and generate more ROS when compared to SS neutrophils

We next investigated whether EG neutrophils displayed functional differences that could, at least in part, contribute to the elevated survival demonstrated by EG larvae. We utilized a method we previously developed to measure neutrophil bacterial killing capacity in larval zebrafish through live time-lapse confocal imaging of *Sal*-GFP-laden neutrophils and measuring the change in intracellular bacterial volume (as detected by *Sal*-GFP fluorescence) over time (‘killing rate’ = Δμm^3^/time) (Astin et al., 2017b). SS and EG larvae were infected with 1,500 CFU *Sal*-GFP at 4 dpf into the hindbrain ventricle and imaged from 2–4 hpsi. This analysis revealed that EG neutrophils demonstrated significantly higher killing rates when compared to SS neutrophils (Figure 1I and Figure S1J). To assess whether this enhanced bactericidal activity of EG neutrophils was heterologous in nature, we next measured the killing rates of SS and EG neutrophils when challenged with another pathogen, GFP-expressing Grampositive *Streptococcus iniae* (*S. iniae*). Similar to their response to Gram-negative *Sal*-GFP, EG neutrophils killed intracellular *S. iniae* faster than SS neutrophils (Figure S1K and L). We then investigated whether EG neutrophils produce more ROS when compared to SS neutrophils by co-injecting the ROS-sensitive fluorescent probe CellROX with *Sal*-GFP at 4 dpf (Figure 1J), as previously described (Astin et al., 2017). This analysis revealed that EG neutrophils produced significantly higher levels of ROS, as detected by CellROX fluorescence intensity (Figure 1K and Figure S1M).

### The enhanced bactericidal activity of EG neutrophils is cell autonomous

When live imaging neutrophil bactericidal activity, we could not discount the possibility that some cells we classified as EG neutrophils may be the few remaining neutrophils we sometimes observed when scoring for neutropenia at 1 dpi (that could have been generated under SS conditions). To enrich for EG neutrophils, we utilized the *Tg(mpx:Dendra2)* neutrophil reporter line (Yoo and Huttenlocher, 2011) to specifically photoconvert neutrophils within the CHT following EG and exclusively measure their killing rates following infection. A 405 nm laser was used to photoconvert Dendra2-expressing neutrophils from green to red within the CHT of SS and EG larvae prior to infection (Figure 2A and B). Photoconverted EG neutrophils had significantly elevated bacterial killing rates when compared to photoconverted SS neutrophils, supporting our earlier results (Figure 2C and Figure S2A).

**Figure 2.**
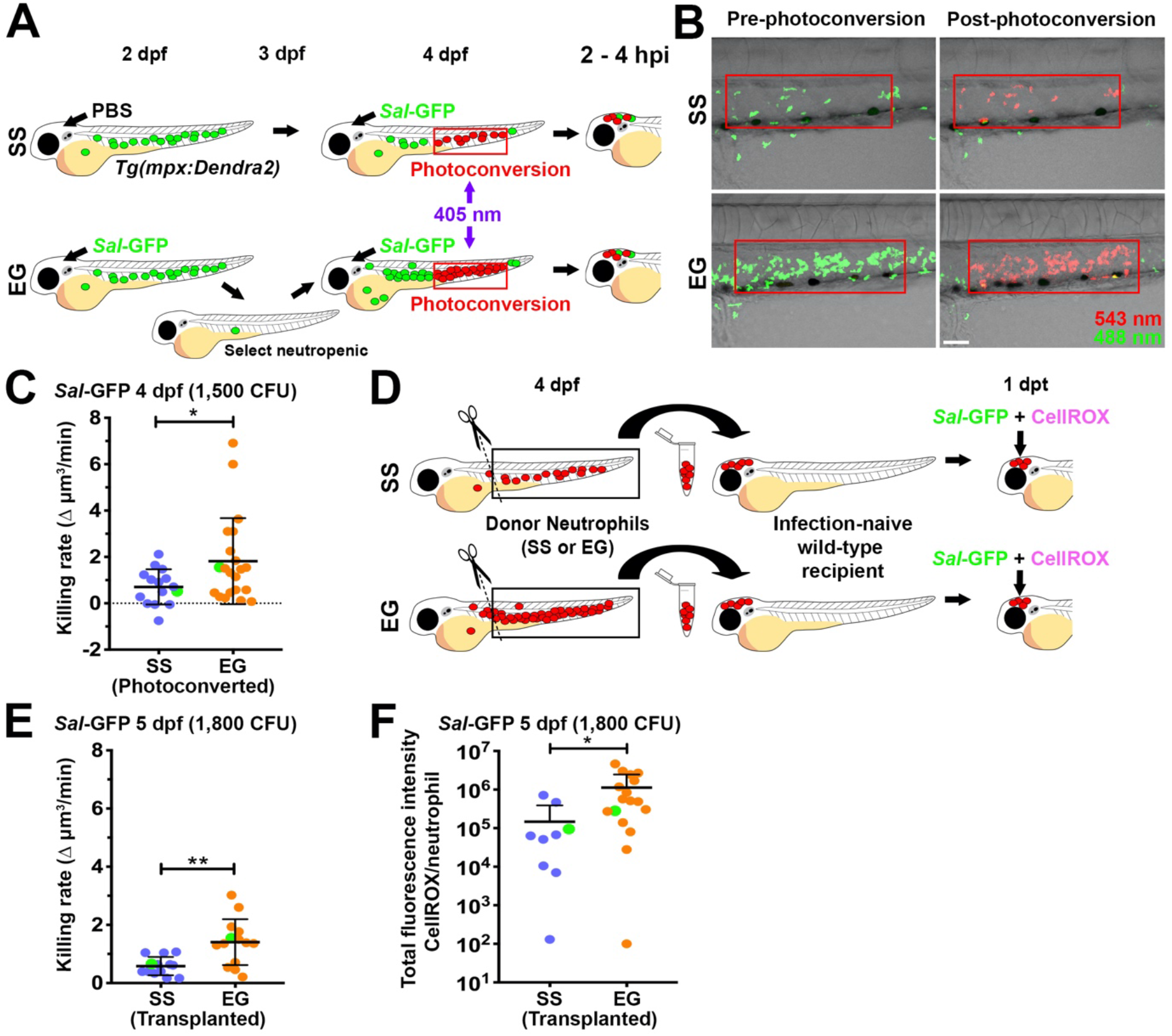
Elevated bactericidal activity and ROS production in EG neutrophils is cell autonomous. (A) Schematic illustrating strategy to enhance selection of EG neutrophils by photoconverting neutrophils within the CHT region. (B) Live imaging of SS and EG neutrophils within the CHT regions of *Tg(mpx:Dendra2*) larvae immediately prior to, and following, photoconversion. Red box marks photoconverted region. (C) Bacterial killing rates of photoconverted SS and EG neutrophils following *Sal*-GFP infection. Green data points highlight killing rates of neutrophils as shown in Figure S2A. (D) Schematic illustrating transplantation of FACS-isolated SS and EG neutrophils into infection-naïve recipient larvae. (E) Bacterial killing rates of transplanted SS and EG neutrophils following *Sal*-GFP infection. Green data points highlight killing rates of neutrophils as shown in Figure S2B. (F) Quantification of ROS production within *Sal*-GFP-laden transplanted SS and EG neutrophils, as detected by CellROX fluorescence. Green data points highlight ROS production of neutrophils as shown in Figure S2C. Error bars, mean ± SD; * p<0.05, **p<0.01; unpaired Student’s t-test (C, E and F). CFU, colony-forming units. Scale bar 50 μm.

We next assessed whether the elevated bacterial killing capacity and ROS production of EG neutrophils was cell autonomous, or the result of being within a previously infected host. Exploiting a neutrophil transplantation protocol we previously developed (Darroch et al., 2020), we examined whether the observed elevated killing rates and ROS production of EG neutrophils was maintained when transplanted into an infection-naïve host. SS and EG neutrophils were isolated from the dissected trunks (containing the AGM and CHT) of SS and EG larvae, respectively, and injected into the hindbrain ventricle of age-matched, infection-naïve recipients (Figure 2D). At 1 day post transplant (dpt), recipients were infected with 1,800 CFU *Sal*-GFP and the bacterial killing rates and ROS production within individual transplanted SS or EG neutrophils were quantified. Transplanted EG neutrophils maintained significantly elevated bacterial killing rates and ROS production when compared to transplanted SS neutrophils (Figure 2E, F and Figure S2B, C).

These results strongly suggest that the enhanced bactericidal activity of EG neutrophils is cell autonomous.

### EG neutrophils have elevated expression of mitochondria-associated genes following infection and utilize mtROS for their enhanced bactericidal activity

Given our neutrophil transplantation experiments strongly suggested that the enhanced capacity of EG neutrophils to kill intracellular bacteria was cell autonomous, we predicted that transcriptional changes may underpin this augmented activity. Bulk RNA-sequencing (RNA-seq) was performed on SS and EG neutrophils before (SS_Before_ and EG_Before_) and after (SS_After_ and EG_After_) infection (Figure 3A). Differentially expressed genes (DEGs) were then identified from pairwise comparisons (Figure 3B). We focused on DEGs that were upregulated in EG_After_ neutrophils when compared to SS_After_ neutrophils as these groups were a direct representation of the neutrophils we functionally characterized in our prior experiments. More specifically, we focused on the 657 genes that were exclusively upregulated after infection in EG_After_ neutrophils when compared to SS_After_ neutrophils (Figure 3B). Gene ontology enrichment analysis focusing on biological processes (GO:BP) performed on this gene set revealed that mitochondrial gene expression and mitochondrial respiratory complex assembly were significantly enriched GO terms (Figure 3C-E). Genes that contributed to the mitochondrial respiratory complex assembly pathway included subunits and assembly factors of electron transport chain (ETC) complex 1 (*ndufs3, ndufs2, ndufs6, nubpl,* and *ndufaf4*) and complex III (*uqcc2, ttc19*) (Figure 3E). Of note, ETC complex I and III are specifically involved in mitochondrial ROS (mtROS) generation (Jastroch et al., 2010). Furthermore, mitochondria have been shown to provide a source of bactericidal ROS to kill intracellular bacteria within phagocytic cells (Abuaita et al., 2018; Dunham-Snary et al., 2022; Geng et al., 2015; Hall et al., 2013; West et al., 2011).

**Figure 3.**
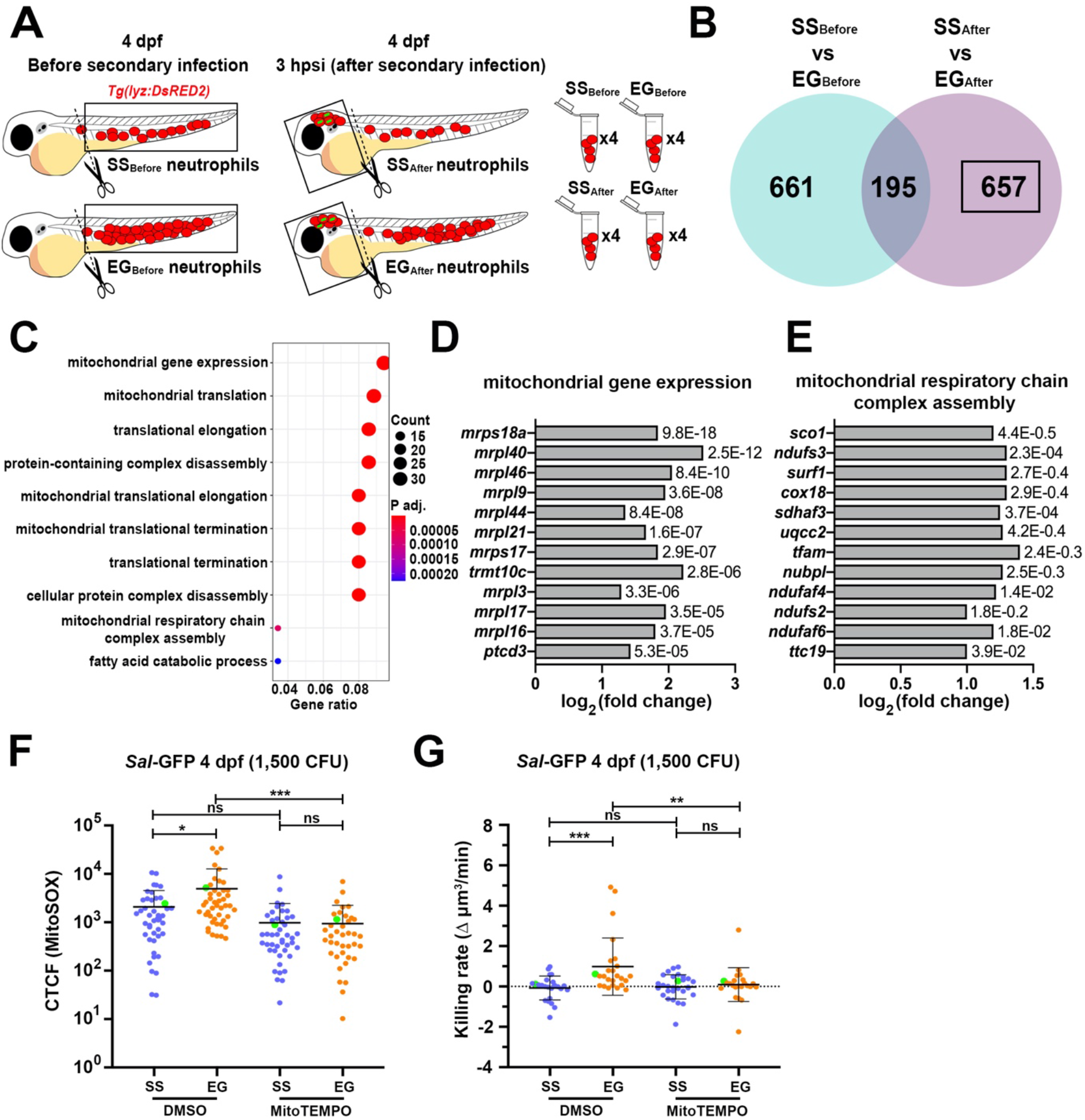
EG neutrophils enhance expression of mitochondria-associated genes following infection and utilize mtROS for their enhanced bactericidal activity. (A) Schematic illustrating sample collection for RNA-sequencing of SS and EG neutrophils before (cells harvested from dissected trunks), and after (cells harvested from dissected heads), infection. (B) Venn diagram showing the number of differentially expressed genes (DEGs) upregulated in EG neutrophils that were unique to the ‘SS_Before_ vs EG_Before_’ and ‘SS_After_ vs EG_After_’ pairwise comparisons and those common to both comparisons. Black box highlights DEGs of interest. (C) GO BP analysis of DEGs of interest. (D) Log2 fold change of genes associated with the ‘mitochondrial gene expression’ GO term. (E) Log2 fold change of genes associated with the ‘mitochondrial respiratory chain complex assembly’ GO term. (F) Quantification of mtROS within *Sal*-GFP laden SS and EG neutrophils in the presence of MitoTEMPO or DMSO (control), as detected by MitoSOX fluorescence. Green data points highlight mtROS within neutrophils as shown in Figure S3A. (G) Bacterial killing rates of SS and EG neutrophils in the presence MitoTEMPO or DMSO (control) following *Sal*-GFP infection. Green data points highlight killing rates of neutrophils as shown in Figure S3B. Error bars, mean ± SD; ns, not significant, * p<0.05, **p<0.01, ***p<0.001; one-way ANOVA with Tukey’s multiple comparisons test (F and G). CFU, colony-forming units.

To investigate if mtROS was contributing to the increased bacterial killing capacity of EG neutrophils, SS and EG *Tg(lyz:EGFP*) larvae were co-injected with 1,500 CFU *Sal*-GFP and the mitochondria-specific fluorescent superoxide indicator MitoSOX Red at 4 dpf. Quantification of mtROS within *Sal*-GFP-laden SS and EG neutrophils revealed that EG neutrophils produced significantly greater amounts of mtROS when compared to SS neutrophils (Figure 3F and Figure S3A). Importantly, this increase in mtROS was abolished in the presence of the mitochondria-targeted antioxidant MitoTEMPO (Figure 3F and Figure S3A). Supporting the bactericidal activity of this mtROS, the increased bacterial killing capacity of EG neutrophils was not observed following MitoTEMPO treatment (Figure 3G and Figure S3B). Together, these data suggest that the increased ability of EG neutrophils to kill intracellular bacteria is, at least in part, due to elevated mtROS production.

### Infection-experienced and *cebpb*-overexpressing HSPCs generate neutrophils with elevated bactericidal activity

Given demand-adapted hematopoiesis is known to be instructed at the level of HSPCs (Baldridge et al., 2010; Baldridge et al., 2011; Essers et al., 2009; Nagai et al., 2006; Takizawa et al., 2012), we speculated that the enhanced bactericidal activity of EG neutrophils was the result of infection-induced changes within HSPCs. Furthermore, with C/EBPβ playing a central role during EG (Akagi et al., 2008; Hirai et al., 2006; Sato et al., 2020) and trained immunity (de Laval et al., 2020) we also explored if overexpressing *cebpb* in HSPCs was sufficient to generate neutrophils with elevated bactericidal activity.

We developed an HSPC transplantation protocol to live image neutrophils derived from transplanted infection-experienced or -naïve HSPCs. To label HSPCs we employed the mouse *Runx1* +24 enhancer to re-created the *Tg(Mmu.Runx1:EGFP*) reporter line previously used to live image and quantify HSPCs (Ng et al., 2010; Zhang et al., 2015), herein referred to as *Tg(Runx1:EGFP*). Of note, the *Tg(Runx1:EGFP*) line also possesses the lens-marking *cry:GFP* transgenesis marker. To validate that our *Tg(Runx1:EGFP*) line marked HSPCs, live confocal imaging revealed that GFP-positive cells resided within the AGM and CHT, were small, spherical and demonstrated ‘endothelial cuddling’, a unique interaction between HSPCs and endothelial cells that was first characterized within larval zebrafish (Figure 4A and Figure S4A) (Tamplin et al., 2015). Additionally, examination of FACS-isolated GFP-positive cells from the dissected trunks of *Tg(Runx1:EGFP*) larvae showed that they resembled HSPCs with large nuclei and sparse ungranulated cytoplasm (Stachura and Traver, 2016) and that they expressed the classical HSPC markers *runx1* and *cmyb* (Kalev-Zylinska et al., 2002; Zhang et al., 2011) (Figure S4B and C). Consistent with our earlier work (Hall et al., 2012), HSPC abundance within *Tg(Runx1:EGFP*) larvae was significantly increased at 1 dpi following injection of *Sal*-GFP into both the hindbrain ventricle and the circulation (~38% and 35% increases for hindbrain and circulation, respectively) (Figure 4A and B).

**Figure 4.**
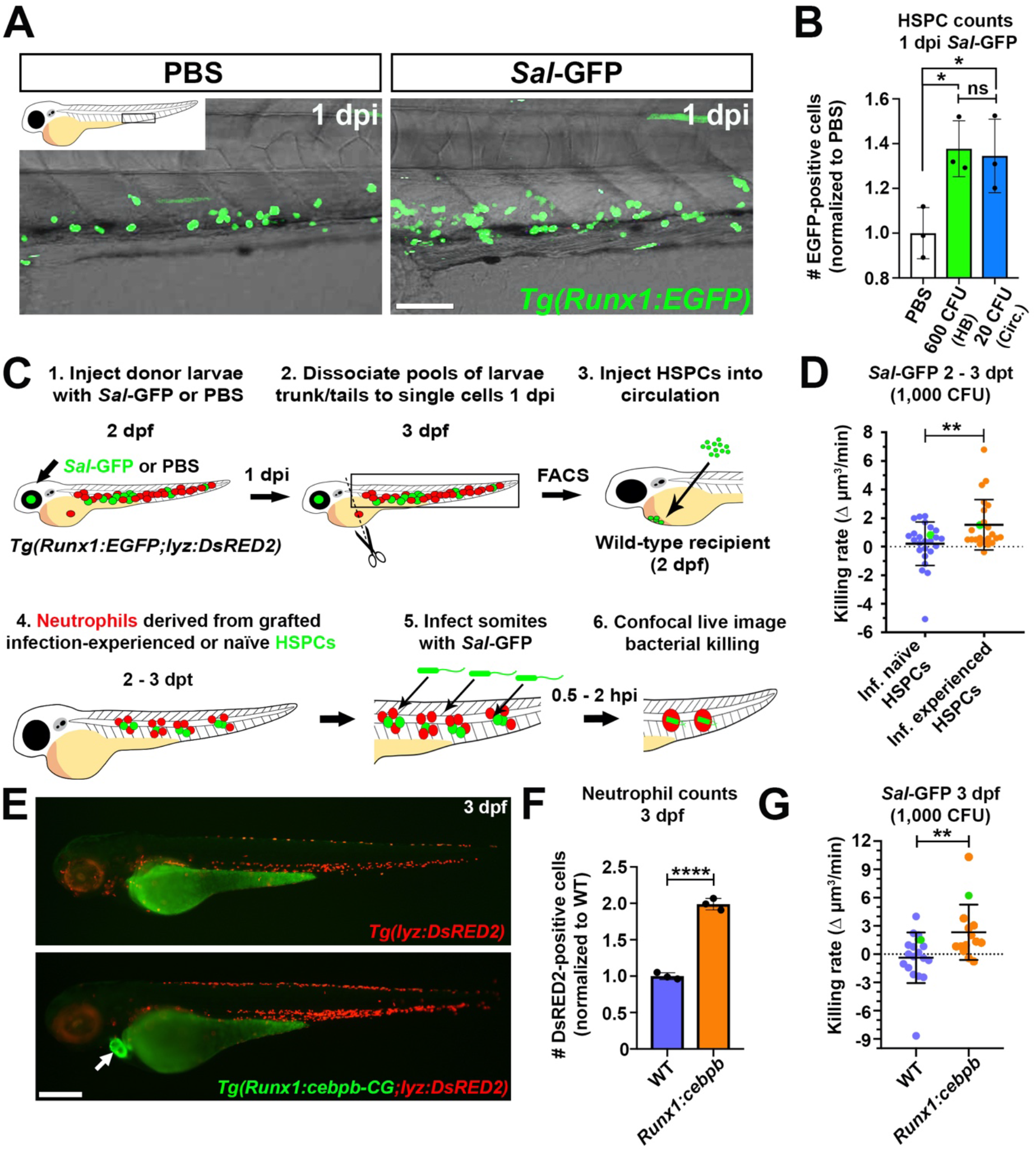
Infection-experienced and *cebpb*-overexpressing HSPCs generate neutrophils with elevated bactericidal activity. (A) Live confocal imaging of HSPCs in the CHT of 3 dpf *Tg(Runx1:EGFP*) larvae, 1 day following injection of PBS or *Sal*-GFP. (B) Flow quantification of HSPCs from dissected trunks of *Tg(Runx1:EGFP*) larvae 1 day following injection with PBS or *Sal*-GFP into the hindbrain ventricle (HB) or circulation (Circ.), n=20 larvae/sample in biological triplicate. (C) Schematic illustrating transplantation strategy to live image neutrophils derived from infection-naïve and -experienced HSPCs. (D) Bacterial killing rates of neutrophils derived from infection-naïve or -experienced HSPCs. Green data points highlight killing rates of neutrophils as shown in Figure S4H. (E) Live imaging of *Tg(lyz:DsRED2*) and *Tg(Runx1:cebpb-CG2;lyzDsRED2*) larvae at 3 dpf. (F) Flow quantification of neutrophils from whole 3 dpf *Tg(lyz:DsRED2*) and *Tg(Runx1:cebpb-CG2;lyzDsRED2*) larvae, respectively (n=10 larvae/sample in biological triplicate). (G) Bacterial killing rates of neutrophils within 3 dpf *Tg(lyz:DsRED2*) and *Tg(Runx1:cebpb-CG2;lyzDsRED2*) larvae, respectively. Green data points highlight killing rates of neutrophils as shown in Figure S4L. Error bars, mean ± SD; ns, not significant, * p<0.05, **p<0.01, ****p<0.0001; one-way ANOVA with Tukey’s multiple comparisons test (B), unpaired Student’s t-test (D, F and G). CFU, colony-forming units. Scale bars, 50 μm in A and 250 μm in E.

To observe neutrophils generated from infection-experienced or -naïve HSPCs, HSPCs were FACS-isolated from 3 dpf *Tg(Runx1:EGFP;lyz:DsRED2*) larvae 1 day following injection with 600 CFU *Sal*-GFP (infection-experienced) or PBS (infection-naïve) (Figure 4C). HSPCs were then transplanted into the circulation of 2 dpf infection-naïve wild-type recipients (Figure 4C). To ensure that no residual *Sal*-GFP was injected alongside HSPCs, transplantation mixtures were plated (Figure S4D). Larvae were then screened for successful engraftment from 1 dpt by confocal microscopy. HSPCs homed to the CHT where they were observed to divide and contribute to the neutrophil lineage from 2 dpt (Figure S4E, F and G). Successfully engrafted infection-naïve recipients that possessed DsRED2-positive neutrophils were then infected with 1,000 CFU *Sal*-GFP and neutrophil bacterial killing rates were quantified (Figure 4C). Neutrophils derived from infection-experienced HSPCs had significantly higher bacterial killing rates when compared to neutrophils derived from infection-naïve HSPCs (Figure 4D and Figure S4H).

To investigate if Cebpb was sufficient to drive emergency granulopoiesis within infection naïve larvae, we generated a transgenic line overexpressing *cebpb* within HSPCs using the mouse *Runx1* +23 enhancer (Tamplin et al., 2015) that incorporated the heart-marking *cmlc2:GFP* (CG2) transgenesis marker (Kwan et al., 2007), herein referred to as *Tg(Runx1:cebpb-CG2*) (Figure S4I and J). Examining HSPC abundance within the dissected trunks of *Tg(Runx1:cebpb-CG2;Runx1:EGFP*) larvae revealed unaltered numbers of HSPCs, when compared to *Tg(Runx1:EGFP*) controls at 3 dpf (Figure S4I and K). This was in contrast to neutrophils that demonstrated a 2-fold increase when quantified from whole *Tg(Runx1:cebpb-CG2;lyz:DsRed2*) larvae (Figure 4E and F). Similar to neutrophils derived from infection-experienced HSPCs, those derived from *cebpb*-overexpressing HSPCs possessed elevated bactericidal activity (Figure 4G and Figure S4L).

Collectively, these data confirm that infection-experienced HSPCs generate neutrophils with elevated bactericidal activity and overexpressing *cebpb* in HSPCs is sufficient to enhance granulopoiesis and produce neutrophils with similarly boosted bactericidal activity.

## DISCUSSION

Here we show that larvae populated with EG neutrophils demonstrate enhanced survival to a secondary lethal infection. We reveal that EG neutrophils can kill intracellular bacteria faster and produce more ROS than SS neutrophils, phenotypes that are maintained following transplantation into infection-naïve hosts. Examining the transcriptomes of EG neutrophils we uncovered an infection-induced transcriptional program that likely operates within EG neutrophils to elevate mtROS production that we show contributes towards their enhanced bactericidal activity. Finally, we provide evidence that the elevated bactericidal activity of EG neutrophils is instructed at the level of infection-experienced HSPCs and that directing *cebpb* expression within infection-naïve HSPCs is sufficient to drive this hematopoietic program.

The enhanced survival of EG larvae to subsequent infection, as shown here, has been similarly reported when driving EG with *Shigella flexneri,* where the authors also show an expanded HSPC compartment (Willis et al., 2018), suggesting this host response is not pathogen-specific, but rather a more heterologous protection mechanism. In this study we reveal that EG neutrophils have enhanced bactericidal activity that operates in a cell-autonomous fashion. This appears to provide heterologous protection as EG neutrophils could kill intracellular bacteria that were the same as those used for the initial ‘priming’ challenge (*Salmonella*) or different (*S. iniae*). Given that EG larvae maintained extended survival in the absence of macrophages suggests that enhanced EG neutrophil bactericidal activity contributes, at least in part, to the elevated survival of EG larvae. Although our EG larvae are populated with increased numbers of neutrophils (~2.5 fold, as shown here and (Hall et al., 2012)), we do not believe that this significantly contributes to increased survival in our infection model given that similar numbers of neutrophils were recruited to infection sites within SS and EG larvae.

Our transcriptomic analysis showed that EG neutrophils had elevated expression of genes associated with mitochondrial respiratory chain complex assembly suggesting enhanced oxidative phosphorylation (OXPHOS) capacity. Supporting this finding, a previous study has demonstrated that neutrophils derived from demand-adapted granulopoiesis have enhanced anti-tumor activity with OXPHOS being similarly identified as a strongly enriched pathway (Kalafati et al., 2020). mtROS is generated as a consequence of OXPHOS largely through electron leakage at ETC complex I and III, and genes involved in the assembly of these complexes were highly represented in our analysis (Jastroch et al., 2010). We revealed that bacteria-laden EG neutrophils produced significantly more mtROS, but not in the presence of the mitochondrial superoxide scavenger MitoTEMPO. Furthermore, MitoTEMPO-treated EG neutrophils had significantly reduced bacterial killing capacity when compared to untreated EG neutrophils indicating that production of mtROS contributes to the elevated bactericidal activity of EG neutrophils. mtROS production is known to increase in both macrophages and neutrophils following infection and has been shown to be directly involved with killing intracellular bacterial in macrophages (Abuaita et al., 2018; Geng et al., 2015; Hall et al., 2013; West et al., 2011). More recently, mitochondria have been uncovered as a bactericidal ROS generator in human neutrophils to kill intracellular *Staphylococcus aureus* (Dunham-Snary et al., 2022). Our work suggests that EG neutrophils adopt an infection-responsive transcriptional program to enhance OXPHOS and boost their bactericidal activity through enhanced mtROS production.

Only recently has evidence emerged that neutrophils contribute to trained immunity. β-glucan-treated mice generate neutrophils with an enhanced anti-tumor phenotype and show enhanced expression of genes involved in phagocytosis and ROS metabolic pathways (Kalafati et al., 2020). In another study, neutrophils from BCG-vaccinated humans displayed a heightened capacity to kill *Candida albicans* and were characterized by enhanced expression of genes involved in activation and degranulation, increased glycolytic rate, and elevated ROS production when stimulated *ex vivo* (Moorlag et al., 2020). To the best of our knowledge, our results show for the first time that infection-experienced HSPCs are directly involved in producing neutrophils with elevated antibacterial activity *in vivo*. To better understand the longevity of protection provided by this source of neutrophils, future work will focus on developing new techniques to live image neutrophils derived from transplanted infection-experienced HSPCs within juvenile and adult zebrafish.

In addition to establishing epigenetic memory in HSPCs during trained immunity (de Laval et al., 2020), C/EBPβ is a transcriptional regulator of emergency granulopoiesis where it operates within HSPCs to drive proliferation and direct myeloid commitment (Akagi et al., 2008; Hirai et al., 2006; Sato et al., 2020). We show that directing *cebpb* expression to HSPCs is sufficient to stimulate granulopoiesis and generate neutrophils with enhanced bactericidal activity, but is not sufficient to increase HSPC numbers. This failure to drive HSPC proliferation may be the result of an inability of our transgenic construct to generate the specific C/EBPβ isoform that induces HSPC proliferation, called LIP (Sato et al., 2020). Performing ChIPseq on infection-experienced and *cebpb*-overexpressing HSPCs will be necessary to reveal if the transcriptional changes we uncovered in EG neutrophils are instructed by epigenetic reprogramming at the level of HSPCs.

In summary, our work suggests that infection-experienced HSPCs contribute to trained immunity by providing a source of ‘demand-adapted’ neutrophils with enhanced bactericidal activity through mtROS production.

## ACKNOWLEDGMENTS

We would like to thank Feng Liu and Len Zon for generously providing the mouse *Runx1* +24 and *Runx1* +23 enhancer constructs, respectively, and Omid Delfi for providing the p3E-*p2a-EGFP* construct used in this study. We thank Leah Rolland for technical support, Soh Kar Yan and Thomas Proft for technical support regarding bacterial work, Stephen Edgar for assistance with flow cytometry, Jacqui Ross for support with confocal microscopy at the Biomedical Imaging Research Unit at the University of Auckland, and Alhad Mahagaonkar and Shantanu Patke for management/maintenance of the zebrafish facility. This study was supported by grants from the Health Research Council of New Zealand (Ref IDs 17/294 and 21/310) and the Royal Society of New Zealand Marsden Fund (Ref ID 20-UOA-183) awarded to C.J.H and by the Ministry of Business, Innovation and Employment of New Zealand awarded to PSC.

## AUTHOR CONTRIBUTIONS

Conceptualization, H.D. and C.J.H.

Formal analysis, H.D.

Funding acquisition, P.S.C. and C.J.H.

Investigation, H.D., P.K., Y.J.S., A.P. and C.J.H.

Methodology, H.D. and C.J.H.

Project administration, J.W.A. and C.J.H.

Resources, P.S.C., J.W.A. and C.J.H.

Supervision, J.W.A. and C.J.H.

Visualization, H.D. and C.J.H.

Writing-original draft, H.D.

Writing-review & editing, H.D, P.S.C., J.W.A. and C.J.H.

## STAR METHDOS

### KEY RESOURCES TABLE

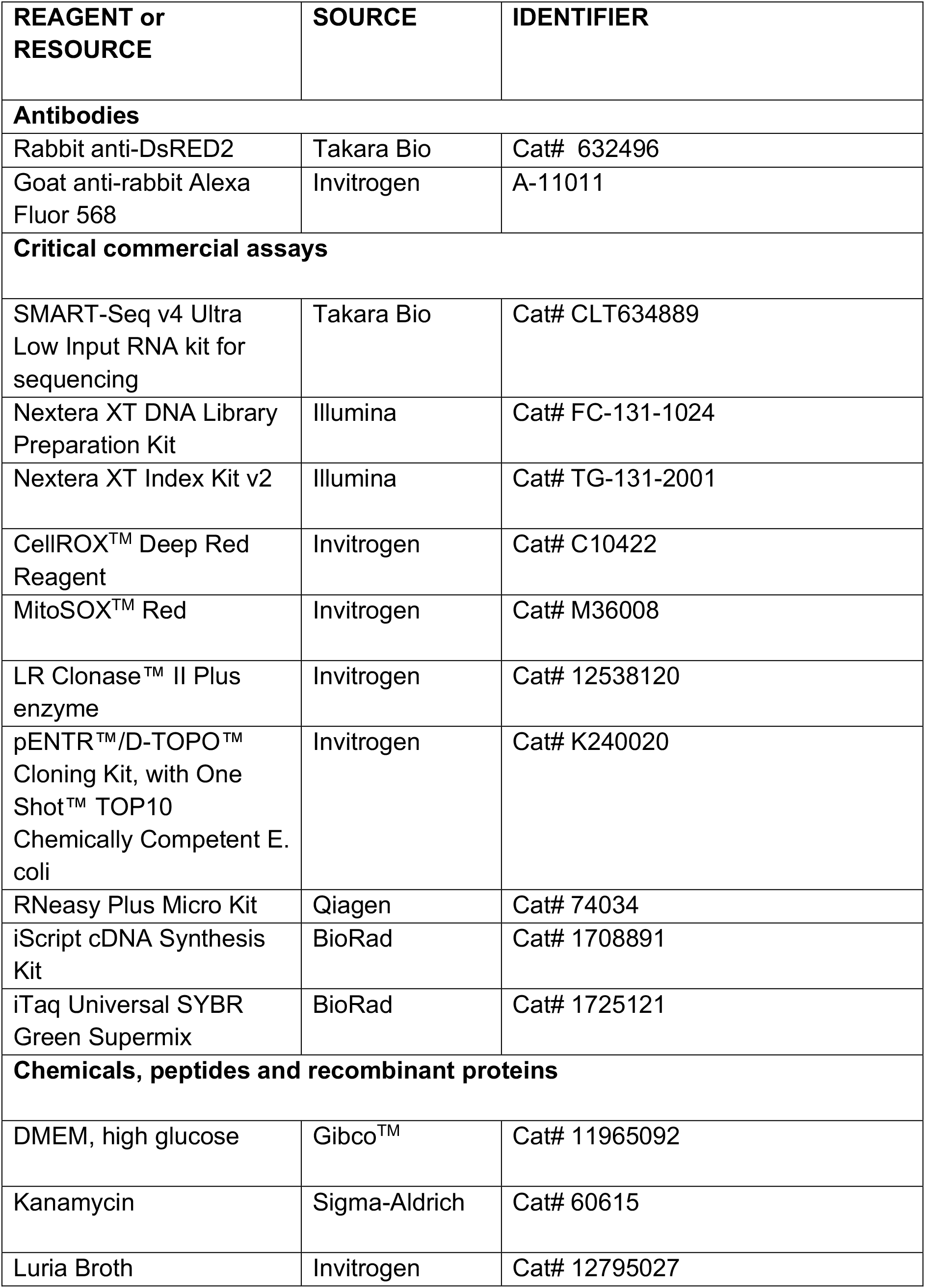
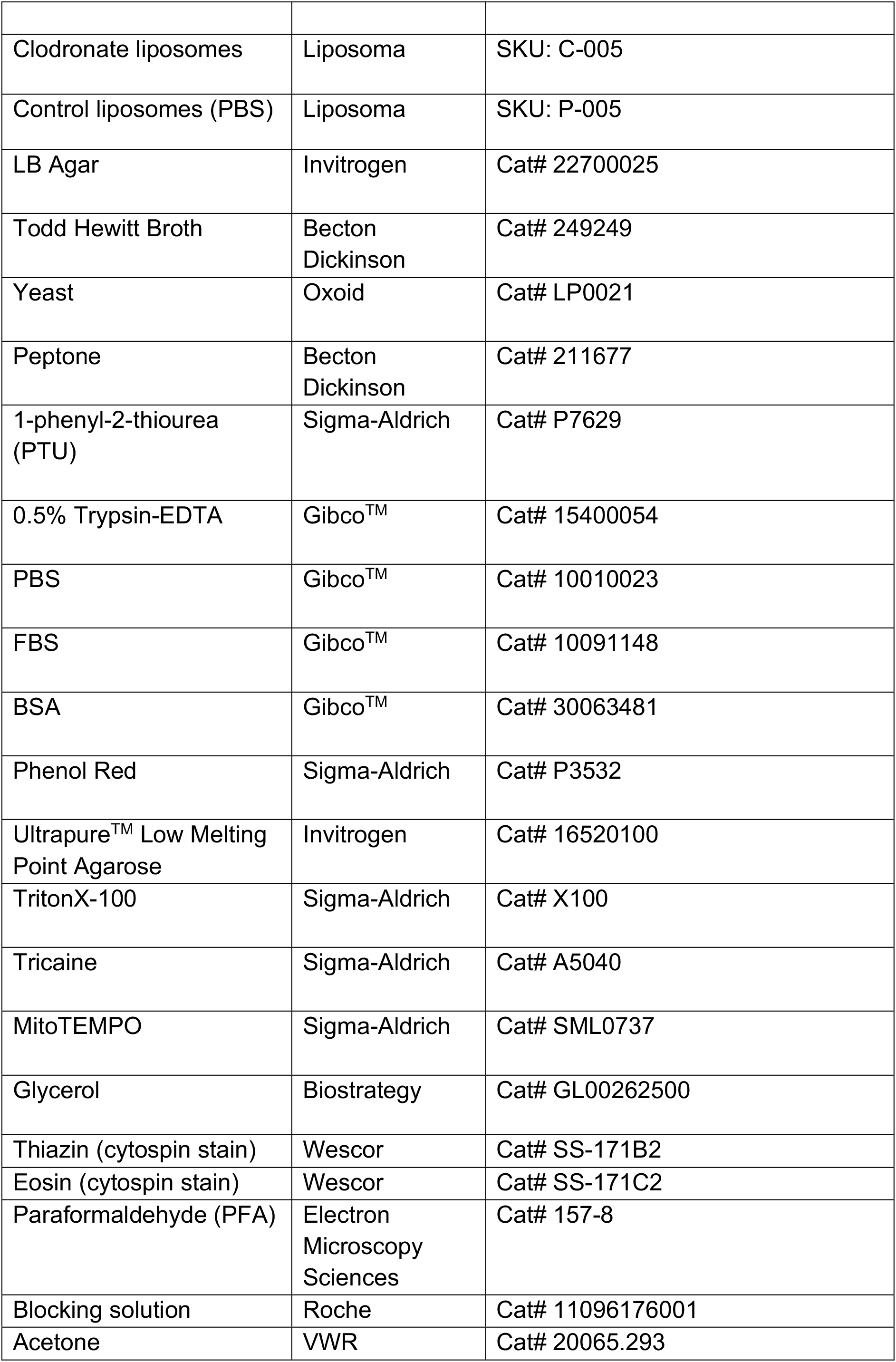
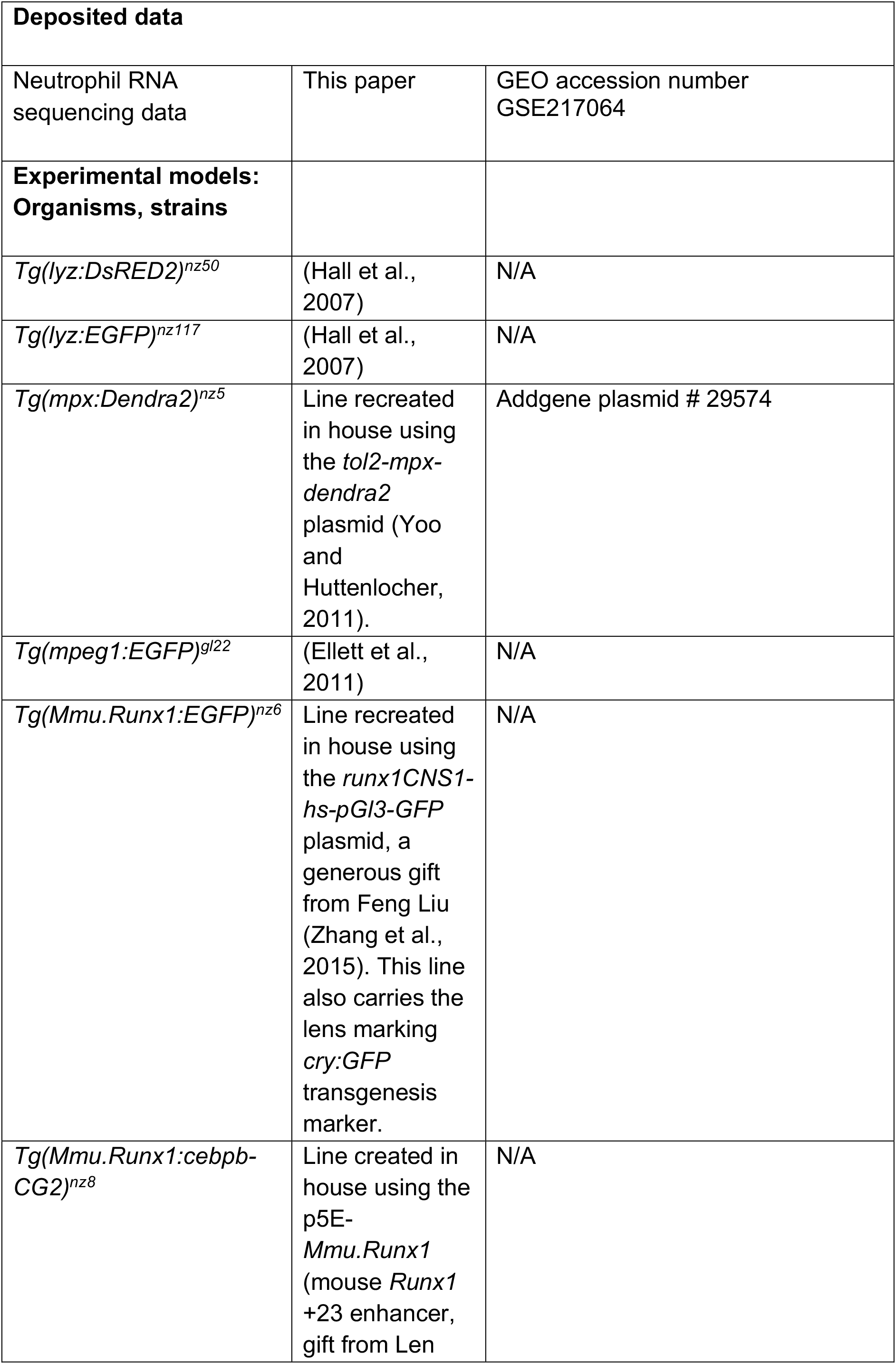
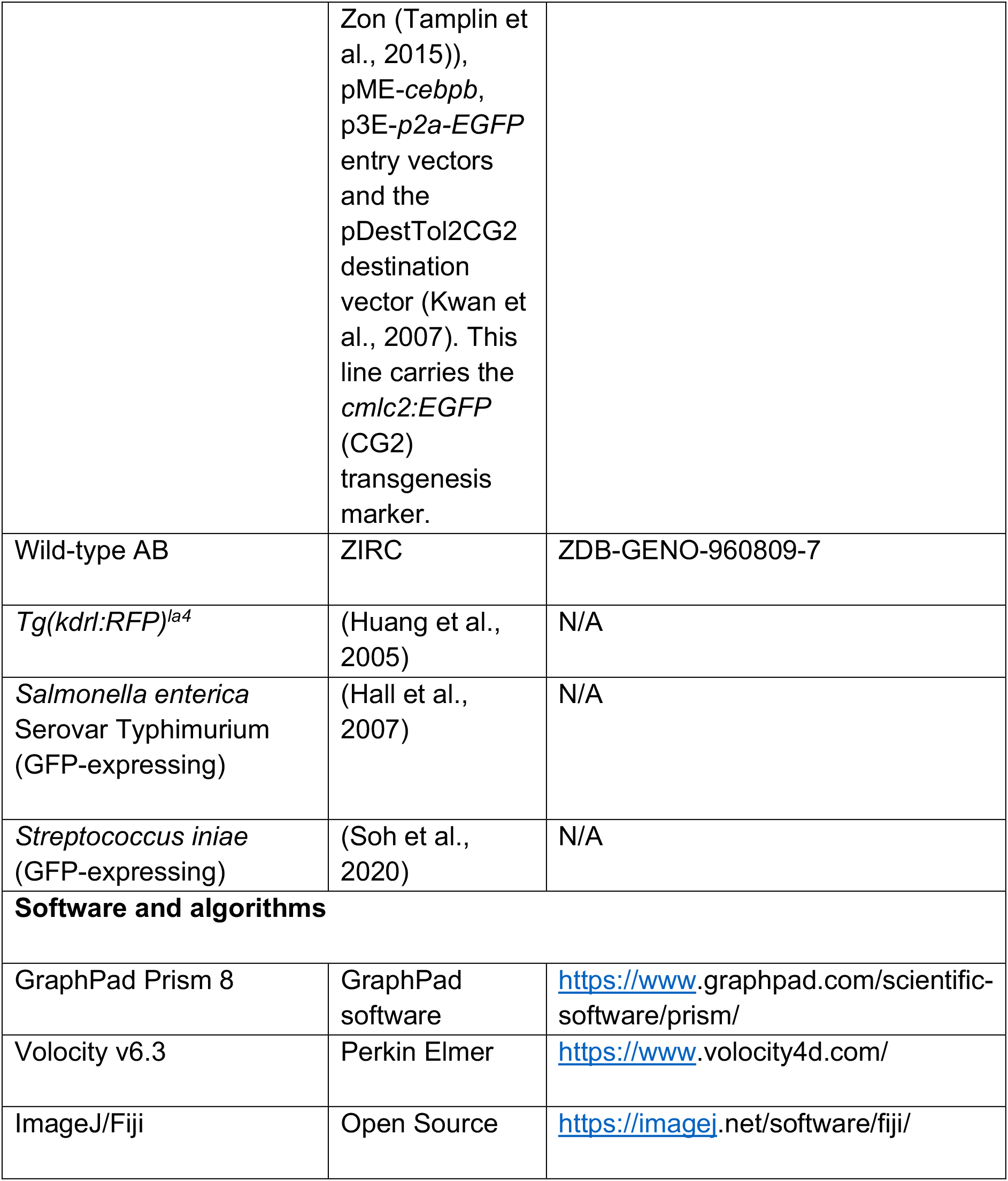

### RESOURCE AVAILABILITY

#### Lead contact

Further information and requests for resources and reagents should be directed to and will be fulfilled by the lead contact, Christopher J. Hall

#### Material availability

All material generated in this study will are available from the lead contact with a material transfer agreement.

#### Experimental model and subject details

##### Zebrafish

All zebrafish research was conducted with the approval of the University of Auckland Animal Ethics Committee (approval numbers AEC001911 and AEC22563).

Adult zebrafish (*Danio rerio*) were sourced from the University of Auckland’s Zebrafish Facility. The facility was on an automatic 14-hour light/10-hour dark light cycle. Larval zebrafish were generated by natural spawning and raised at 28°C in E3 medium supplemented with 0.003% phenylthiourea (PTU) to inhibit the development of pigmentation. Where appropriate, zebrafish were anesthetized by supplementing E3 with 4% (v/v) tricaine in E3 medium. Wild-type AB zebrafish were obtained from the Zebrafish International Resource Centre (ZIRC). *Tg(lyz:DsRED2)^nz50^* and *Tg(lyz:EGFP)^nz117^* neutrophil-specific reporter lines were utilized in this study, as well *Tg(mpeg1:EGFP)^gl22^* and *Tg(kdrl:RFP)^la4^* that label macrophages and blood vessels, respectively (Ellett et al., 2011; Hall et al., 2007; Huang et al., 2005). In addition, the *Tg(mpx:Dendra2)^nz5^* and *Tg(Mmu.Runx1:EGFP)^nz6^* (herein referred to as *Tg(Runx1:EGFP*)) reporter lines were re-created in-house from gifted plasmids for use in this study that mark neutrophils and hematopoietic stem and progenitor cells (HSPCs), respectively (Yoo and Huttenlocher, 2011; Zhang et al., 2015).

##### Bacterial cultures

The following bacterial strains were used for in vivo studies: *Salmonella enterica* serovar Typhimurium and *Streptococcus iniae,* that both express GFP.

### METHOD DETAILS

#### Generation of the *Tg(Mmu.Runx1:cebpb-CG2*) reporter line

The *Tg(Mmu.Runx1:cebpb-CG2*) line (herein referred to as *Tg(Runx1:cebpb-CG2)*) was generated via Gateway cloning (Invitrogen). The zebrafish orthologue of C/EBPβ (encoded by *cebpb*) was ordered as a gBlock® Gene Fragment from integrated DNA technologies (IDT). This included the 840 bp sequence (stop codon removed) of the single *cebpb* exon (ENSDARG00000042725) with an additional 4 bp sequence (CACC) at the 5’ end of the gBlock® to facilitate directional cloning. This sequence was cloned into the pENTR/D-TOPO vector as per the manufacturer’s protocol (Invitrogen) to produce the pME-*cebpb* plasmid. pME-*cebpb*, *p5E-Mmu.Runx1* (mouse *Runx1* +23 enhancer, gift from Len Zon (Tamplin et al., 2015)) and p3E-*p2a-EGFP* (a gift from Omid Delfi) were cloned into the pDestTol2CG2 destination vector (Kwan et al., 2007) via LR clonase II plus recombination, as per the manufacturer’s protocol (Invitrogen). pDESTTol2CG2 contains the heart-marking *cmlc2:EGFP* transgenesis marker. 1 nL of this construct was injected into single-cell stage AB embryos with the following injection mix: 1 μL (200 ng) plasmid DNA, 1 μL (250 ng) transposase mRNA, 3 μL phenol red, 5 μL ultrapure water. At 24 hours post-injection, embryos were selected for the green heart marker, raised to sexual maturity and outcrossed to AB to identify germline transmitting founders. Positive F1 progeny were then raised as a stable transgenic line.

#### Bacterial injections

GFP-expressing *Salmonella enterica* (herein referred to as *Sal*-GFP) was prepared by inoculating 4 mL of Luria Bertani (LB) broth supplemented with 25 μg/mL kanamycin and culturing overnight at 28°C. This was diluted 1:10 in equal parts LB and DMEM. This subculture was then incubated at 37°C for 45 min whilst shaking at 200 revolutions per min (rpm). The bacterial pellet was obtained by centrifugation and resuspended with sterile PBS and 0.25% phenol red to the desired final concentration.

*Streptococcus iniae (S. iniae*) was prepared by inoculating 2 mL of Todd Hewitt broth with 2% (w/v) yeast and 20% (w/v) peptone (THY+P) supplemented with 50 μg/mL kanamycin and cultured for 24 h at 37°C. This was diluted 1:10 in THY+P broth and incubated at 37°C without shaking until the OD_600nm_ was 0.4. 1 mL of the culture was centrifuged and resuspended in 1 mL of THY+P broth supplemented with 25% glycerol and stored at −80°C until required. Immediately prior to injecting, frozen stocks were thawed, centrifuged, and resuspended in sterile PBS containing 0.25% phenol red to the desired final concentration.

Larvae were raised to the appropriate developmental time point, anesthetized in 4% tricaine, and then arrayed laterally in 3% (w/v) methylcellulose in E3 medium. Approximately 1 nL of the bacterial injection mix was microinjected into the appropriate anatomical location (hindbrain ventricle, circulation or somite), as previously described (Hall et al., 2012). The dose of bacteria was validated for each experiment by diluting an injection bolus 1:10 and 1:100 in sterile PBS, both before and after the infections, and plating on LB agar (*Sal*-GFP) or THY+P agar (*S. iniae*) supplemented with kanamycin.

#### Survival analysis

Survival analysis was performed as previously described (Hall et al., 2013). Larvae were monitored for 5 days following infection and dead larvae, as judged by cardiac arrest, were removed and counted at 24 h intervals.

#### Bacterial CFU enumeration

Bacterial CFUs within individual larvae was performed as previously described (Hall et al., 2013). In brief, larvae were rinsed in E3 medium at set timepoints following infection and placed into separate microtubes containing 100 μL of sterile PBS supplemented with 1% Triton X-100. Each larva was homogenized with a sterile pestle and the homogenate was diluted in a 100-fold dilution series. 10 μL of each dilution (max 1:10^8^) was spot plated in triplicate onto LB agar supplemented with 25 μg/mL kanamycin. Plates were incubated at 28°C overnight. Bacterial burdens were determined by calculating the average number of colonies recovered at the highest dilution that grew colonies in each replicate. Bacterial burdens were quantified for 8 to 15 larvae per time point.

#### Confocal microscopy

Larvae were live imaged on an Olympus Fluoview FV1000 laser scanning confocal microscope equipped with an incubation chamber set to 29°C. Larvae were anesthetized with 4% tricaine and mounted in 0.7-1% (w/v) low melting point agarose in E3 medium supplemented with 0.003% PTU and 3.2% tricaine. The lower (0.7%) concentration of agarose was used for longer term time-lapse imaging. Images were captured with a 60X water immersion lens, a scan format of 512 x 512 pixels and a zoom setting of 1.5-2.5X. All imaging parameters were kept identical within, and across, imaging experiments when measuring neutrophil killing rates and quantifying CellROX and MitoSOX fluorescence within bacteria-laden neutrophils, including laser voltage, resolution, scanning speed, offset, and gain.

#### Macrophage ablation

For liposome clodronate-mediated macrophage ablation, liposomes were injected into both the hindbrain ventricle and circulation as previously described (Hall et al., 2018). Of note, given the requirement of blood circulation for recruitment of innate immune cells to the hindbrain infection site, only liposome-injected larvae with unaffected blood flow were used for subsequent experiments.

#### Immunofluorescence

Immunofluorescent detection of neutrophils within *Tg(lyz:DsRED2*) larvae was performed as previously described (Hall et al., 2013). The primary and secondary antibodies used for the detection of DsRED2 their dilutions were: rabbit anti-DsRED2, 1:400 (Takara Bio; cat# 632496) and goat anti-rabbit Alexa Fluor 568, 1:400 (Invitrogen, A-11011).

#### Quantifying neutrophil killing rates, ROS and mtROS production

Neutrophil bacterial killing rates were measured at 2-4 hours post infected by timelapse confocal microscopy as previously described (Astin et al., 2017). Z-stacks (1.5 μm step size) of individual bacteria-laden neutrophils were collected for 8-15 min. Imaged neutrophils had to satisfy the following criteria to be included in a dataset: there was no evidence of additional bacterial phagocytosis over the imaging period; the neutrophil needed to remain within the X-, Y- and Z-dimensions throughout the imaging period; and the imaging period needed to be between 8-15 min, which was sufficient to detect a killing rate while minimizing photobleaching. Time-lapse movies were analyzed using Volocity v6.3 image analysis software. Within bacteria-laden neutrophils that satisfied the above criteria, the volume (μm^3^) of GFP-labeled bacteria was measured using the volume measurement tool in Volocity at the beginning and end of each time-lapse movie. First, objects were identified using GFP fluorescence signal intensity (using a constant threshold setting), a region of interest (ROI) was manually positioned around the neutrophil in X-, Y- and Z-dimensions and the sum of the volumes for all objects within the ROI was quantified to give a total volume. From this neutrophil killing rates were calculated as (initial volume – final volume)/time, or Δ μm^3^/min, where a positive value is interpreted as intracellular bacterial killing and a negative value as bacterial growth.

CellROX Deep Red Reagent, a cell-permeable probe that fluoresces following oxidation by ROS, was used to measure ROS production in neutrophils as previously described (Astin et al., 2017). In brief, the bacterial injection mix was supplemented with 50 μM CellROX and delivered into the hindbrain ventricle by microinjection. Z-stacks (1.5 μm step size) of bacteria-laden neutrophils were then imaged by confocal microscopy at 2-4 hours post infection. The total fluorescence intensity of CellROX within neutrophils containing intracellular bacteria was measured using Volocity v6.3. Z-stacks were first manually inspected to select for neutrophils containing bacteria. Next, a region of interest was drawn around individual bacteria-laden neutrophils in the X-, Y- and Z-dimensions. Employing the threshold function (using a constant threshold setting) for the CellROX channel (excitation/emission maxima of 640/665 nm), the sum of the greyscale intensity values of individual voxels within the region of interest were measured and plotted as the total fluorescence intensity CellROX/neutrophil.

MitoSOX Red (excitation/emission maxima of 396/610 nm), a cell-permeable fluorescent probe selective for mitochondria-specific superoxide, was used to measure mitochondrial ROS (mtROS) in neutrophils. To quantify mtROS we adapted a protocol we previously used to measure mtROS within larval zebrafish macrophages using MitoSOX (Hall et al., 2013). In brief, the bacterial injection mix was supplemented with 50 μM MitoSOX. Live confocal imaging was used to take single Z-stacks (1.5 μm step size) of bacteria-laden neutrophils from 2-4 hours post infection. The fluorescence intensity of MitoSOX within neutrophils containing intracellular bacteria was measured using ImageJ/Fiji. Z-stacks were first manually inspected to select for neutrophils containing bacteria. Next, a region of interest was drawn around individual bacteria-laden neutrophils in the X-, Y- and Z-dimensions and the controlled total cell fluorescence (CTCF) of MitoSOX/neutrophil was determined using ImageJ/Fiji as previously described (Elks et al., 2013). CTCF was used for this probe to correct for background noise.

#### Chemical treatments

MitoTEMPO, a mitochondrial superoxide scavenger, was injected immediately following bacterial infection into the hindbrain ventricle at a concentration of 250 μM in DMSO as previously described (Hall et al., 2018).

#### Photoconversion

*Tg(mpx:Dendra2*) larvae were viewed under a 20x water dipping objective lens. A ROI was selected using the FluoView FV1000 software around the Dendra2-expressing cells of interest in the CHT and a 405 nm laser was targeted to the ROI for 1 minute to photoconvert Dendra2 from green (excitation/emission maxima of 490/507 nm) to red (excitation/emission maxima of 553/573 nm) fluorescence.

#### Live fluorescence microscopy

Larvae were anesthetized with 4% tricaine and arrayed in 3% methylcellulose in E3. Images were taken using a DS-U2/L2 camera fitted to a Nikon SMZ1500 fluorescence stereomicroscope.

#### Flow cytometry and FACS

Larvae were dissociated for flow cytometry/FACS as previously described with modifications (Covassin et al., 2006). In brief, dechorionated larvae were rinsed in calcium-free Ringer’s solution supplemented with 2 mM MgCl2 and 10 mM D+ glucose on ice for 15 min. Larvae were de-yolked by repeated passage through a 200 μL pipette tip and the supernatant discarded. Larvae were digested in 0.25% trypsin-EDTA in PBS for 1-1.5 h at 28°C with manual agitation every 10 min. The digestion was inhibited by adding CaCl2 to a final concentration of 1 mM and 5% FBS. The dissociated material was centrifuged at 260 x g for 5 min at 4°C and the supernatant removed by decanting. The cell pellet was resuspended in ice-cold PBS supplemented with 5% FBS. Cells were passed through a 40 μm cell strainer (BD falcon) and stored on ice prior to flow cytometry or FACS. Flow cytometry was performed on a BD LSR II flow cytometer and cells were gated based on forward and side scatter characteristics, and GFP expression. Cells were FACS-isolated on a BD FACS Aria II into PBS supplemented with 5% FBS and either stored on ice prior for transplantation or processed for expression analysis by qPCR or RNA sequencing.

#### Cytospin preparation and cell staining

Cells were collected via FACS as described above. Samples were spun at 200 x g for 5 minutes at 4°C. The supernatant was discarded and the cell pellet resuspended in 250 μL of cytospin buffer (3 mM EDTA, 2% BSA, 1% FBS in 1x PBS). Cells were then fixed to slides using an Aerospray Hematology Pro cytospin, and stained immediately (Wright-Giemsa) using the in-built slide staining mechanism.

#### Cell transplantation

Neutrophils or HSPCs were isolated from donor larvae and transplanted into recipient larvae as previously described with modifications (Darroch et al., 2020). In brief, around 5,000 cells were FACS-isolated into PBS supplemented with 5% FBS and then centrifuged at 260 x g for 10 min at 4°C. The supernatant was removed by pipetting leaving ~20 μL behind to resuspend the pellet. Recipient larvae were anesthetized with 4% tricaine and arrayed in 3% methylcellulose in E3 medium. 3 μL of the cell resuspension was loaded into a microinjection needle and an MPPO-2 Pressure Injector (Apllied Scientific Instrumentation) was used to inject cells into the appropriate location within recipient larvae (hindbrain ventricle for neutrophils, circulation for HSPCs). Transplanted larvae were returned to fresh E3 with 0.003% PTU and monitored for successful engraftment using confocal or fluorescent microscopy from 1 day post transplant.

#### RNA extraction and quantitative PCR (qPCR)

Total RNA was extracted from FACS-isolated sorted cells using the RNeasy Micro Plus Kit (Qiagen) where cells were directly sorted into 350 μL of RLT Plus buffer and RNA extracted as per the manufacturer’s protocol. Total RNA was reverse transcribed into cDNA using the iScript cDNA Synthesis Kit (BioRad) and qPCR was performed using the iTaq Universal SYBR Green Supermix (BioRad) and the QuantStudio 6K Flex Real-Time PCR System (Life Technologies, Thermo Fisher Scientific). The following primer pairs were used for *runx1,* forward 5’-AATGACCTGCGTTTCGTGGG-3’, reverse 5’-TGTCGGTGGCGTCGTGG-3’; *cmyb,* forward 5’-ACAACAGGCACTACCAATCTCC-3’, reverse 5’-CAATGCCAACCGAACTGTCC-3’; *cebpb*, forward 5’-CTTTCCACAGCACTAACGCC-3’, reverse 5’-AGTCTATGGCTTTCTCGTGC-3’; *ef1a,* forward 5’-TGCCTTCGTCCCAATTTCAG-3’, reverse 5’-TACCCTCCTTGCGCTCAATC-3’. Each qPCR experiment was performed in biological and technical triplicate. Expression levels were normalized to *ef1α* and calculated using the ΔΔCt method.

#### RNA-sequencing and data analysis

Larvae were prepared for FACS as described above, with a change in protocol that the cells were sorted into 0.1% BSA in PBS instead of 5% FBS to prevent inhibition of downstream enzymatic reactions. 1,000 cells per sample were sorted directly into a ‘lo-bind’ 1.5 mL Eppendorf tube containing 1 μL 10X lysis buffer, 1 μL Rnase inhibitor and 5 μL nuclease-free water. Nuclease-free water was then added to 10.5 μL. Sorted cells were then vortexed briefly to homogenize and lyse the cells, briefly centrifuged, and then stored at −80°C. Frozen samples were thawed on ice prior to cDNA synthesis as per the SMART-Seq v4 Ultra Low Input RNA kit for sequencing (Takara Bio) manufacturer’s protocol. cDNA from each sample was run on an Agilent 2100 Bioanalyzer to measure cDNA fragment size (in base pairs), concentration, and purity. 1 ng total cDNA was the input for each sample into the Nextera XT DNA Library Preparation Kit (Illumina). DNA libraries were prepared as per the manufacturer’s protocol. Libraries were checked for size distribution, purity, and concentration on an Agilent 2100 Bioanalyser. Libraries were sequenced using NextSeq 500 in single end reads, 75 base pair read length.

Single end reads were checked for quality using FastQC and filtered, cleaned reads were mapped to the zebrafish reference genome (GRCz11) from Ensembl using HISAT2. Expression levels were measured as Fragments Per Kilobase of transcript per Million mapped reads (FPKM) using the Cuffquant and Cuffnorm components of Cufflinks software. DESeq was then used to analyze the differentially expressed genes (DEGs) between sets of two groups as a control-treatment pairwise comparison. The expression fold change (log2FC) was set to >1 and the false discovery rate (FDR) was set to <0.05 as screening criteria for DEGs. DEGs were functionally annotated using gene ontology (GO) enrichment analysis. Analysis of RNA sequencing data was performed with assistance from CD genomics (https://www.cd-genomics.com/).

#### Statistical significance

Statistical significance was determined using GraphPad Prism 8. Multiple comparison analysis was carried out by ordinary one-way ANOVA with Tukey’s multiple comparisons test. Pair-wise comparisons were carried out by Student’s unpaired t-test. Kaplan-Meier survival plot analysis was carried out by the Gehan-Breslow-Wilcoxon method. Statistical significance was defined by p values <0.05. The statistical test used, and p values, are indicated in each figure legend.

#### Data and code availability

Neutrophil RNA sequencing data have been deposited in the NCBI Gene Expression Omnibus (GEO) under accession number GSE217064 and are publicly available upon publication.

Any additional information required to reanalyze the data reported in this paper is available from the lead contact upon request.

