## Supplemental figures for "Infection-experienced HSPCs protect against infections by generating neutrophils with enhanced bactericidal activity"

### Supplementary data

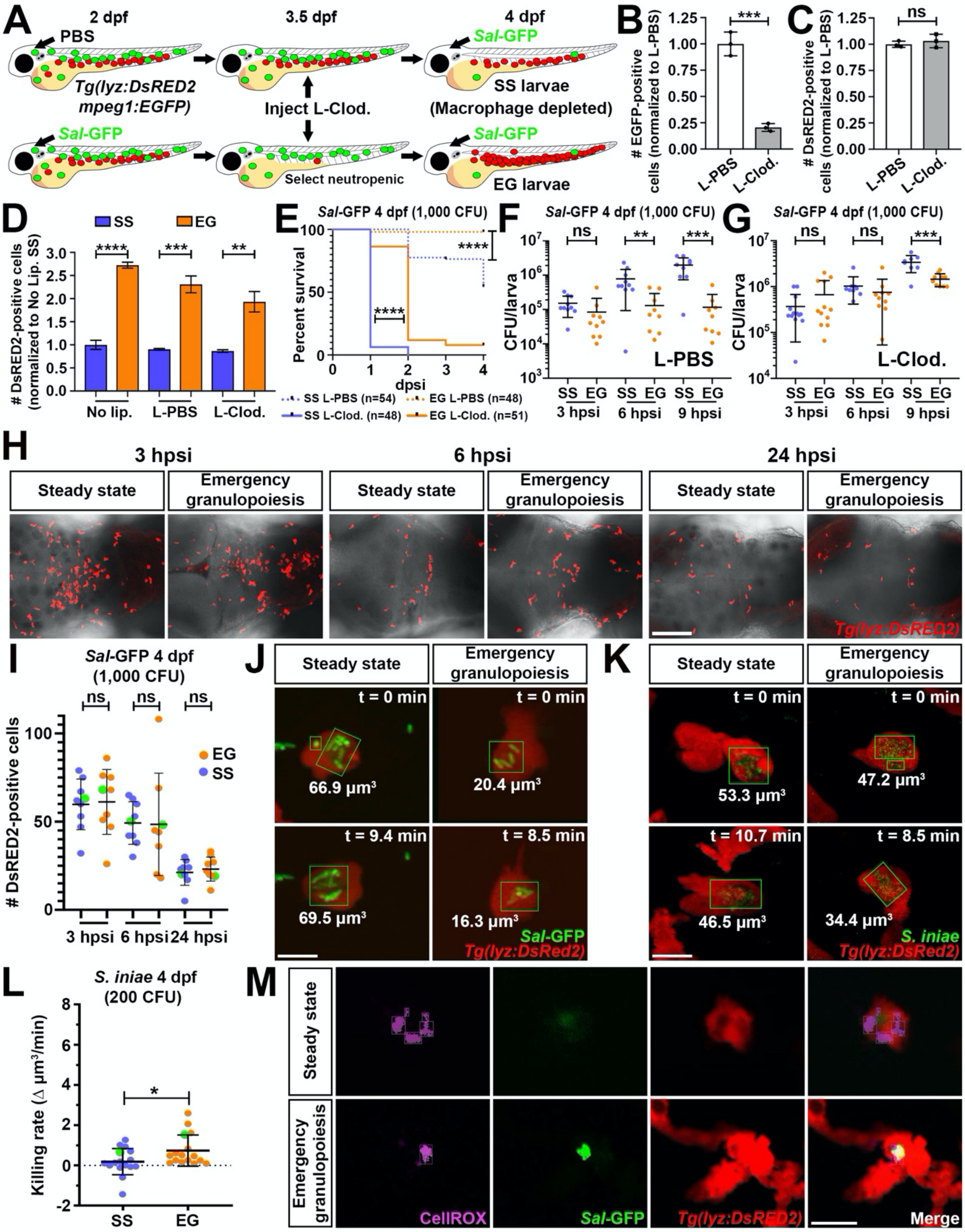

**Figure S1. EG larvae have extended survival to infection in the absence of macrophages, similar numbers of neutrophils are recruited to infections within SS and EG larvae and live confocal imaging of neutrophils following *Sal*-GFP and *S. iniae* infection. Related to Figure 1.**

(A) Schematic illustrating strategy to generate SS and EG larvae depleted of macrophages. *Tg(lyz:DsRED2;mpeg1:EGFP)* larvae were injected with PBS or *Sal*-GFP at 2 dpf. At 3 dpf, neutropenic larvae were selected from the *Sal*-GFP-injected cohort. Liposomal clodronate (L-Clod.) was injected into the hindbrain and circulation of both cohorts at 3.5 dpf. At 4 dpf, macrophage-depleted SS and EG larvae were infected with *Sal*-GFP to determine survival and bacterial burdens. (B and C) Flow quantification of EGFP-expressing macrophages (B) and DsRED2-expressing neutrophils (C) from whole 4 dpf *Tg(lyz:DsRED2;mpeg1:EGFP)* larvae, following L-Clod. injection, compared to L-PBS injection controls (n=20 larvae/sample in biological triplicate). (D) Flow quantification of DsRED2-expressing neutrophils from whole 4 dpf SS and EG *Tg(lyz:DsRED2;mpeg1:EGFP)* larvae, 1 day following L-Clod. injection, compared to L-PBS injection and no liposome (No lip.) controls (n=20 larvae/sample in biological triplicate). (E) Kaplan-Meier graphs showing survival of SS L-PBS, SS L-Clod., EG L-PBS and EG L-Clod. larvae over 4 days post secondary injection (dpsi) with *Sal*-GFP at 4 dpf. (F) Bacterial burdens within individual L-PBS-injected SS and EG larvae at 3, 6 and 9 hpsi with *Sal*-GFP at 4 dpf. (G) Bacterial burdens within individual L-Clod.-injected SS and EG larvae at 3, 6 and 9 hpsi with *Sal*-GFP at 4 dpf. (H) Immunofluorescence detection of neutrophils in the hindbrain ventricles of SS and EG *Tg(lyz:DsRED2)* larvae at 3, 6, and 24 hpsi with *Sal*-GFP. (I) Quantification of neutrophils as detected in H. Green data points highlight larvae shown in H. (J) Frame shots from live time-lapse confocal imaging of SS and EG neutrophils within *Tg(lyz:DsRED2)* larvae showing volumes of intracellular *Sal*-GFP at the beginning (t=0) and end of the time-lapse experiment. (K) Frame shots from live time-lapse confocal imaging of SS and EG neutrophils within *Tg(lyz:DsRED2)* larvae showing volumes of intracellular *S. iniae* at the beginning (t=0) and end of the time-lapse experiment. (L) Bacterial killing rates of SS and EG neutrophils following *S. iniae* infection. Green data points highlight killing rates of neutrophils as shown in K. (M) Live confocal imaging of ROS production within individual *Sal*-GFP-laden SS and EG neutrophils within *Tg(lyz:DsRED2)* larvae, as detected by CellROX fluorescence. Error bars, mean ± SD; ns, not significant, * $p<0.05$, ** $p<0.01$, *** $p<0.001$, **** $p<0.0001$;

unpaired Student’s t-test (B, C, D, F, G, I and L), Gehan-Breslow-Wilcoxon test (E). CFU, colony-forming units. Scale bars, 100 μm in H and 10 μm in J, K and M.

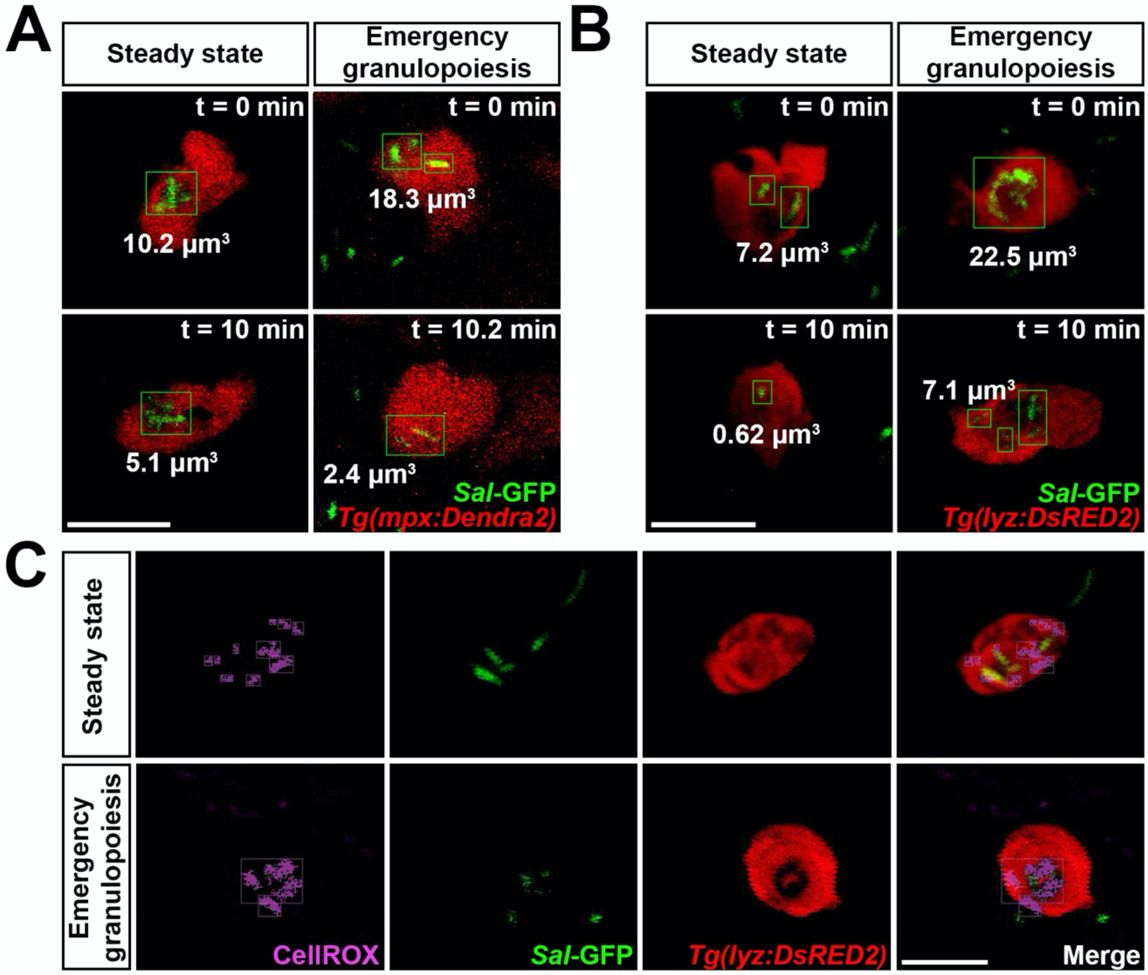


**Figure S2. Confocal imaging of photoconverted and transplanted neutrophils following *Sal*-GFP infection. Related to Figure 2.**

(A) Frame shots from live time-lapse confocal imaging of photoconverted SS and EG neutrophils within *Tg(mpx:Dendra2)* larvae showing volumes of intracellular *Sal*-GFP at the beginning (t=0) and end of the time-lapse experiment. (B) Frame shots from live time-lapse confocal imaging of SS and EG neutrophils transplanted from *Tg(lyz:DsRED2)* larvae into WT recipients, showing volumes of intracellular *Sal*-GFP at the beginning (t=0) and end of the time-lapse experiment. (C) Live confocal imaging of ROS production within individual *Sal*-GFP-laden SS and EG neutrophils transplanted from *Tg(lyz:DsRED2)* larvae into WT recipients, as detected by CellROX fluorescence. CFU, colony-forming units. Scale bar 10 µm.

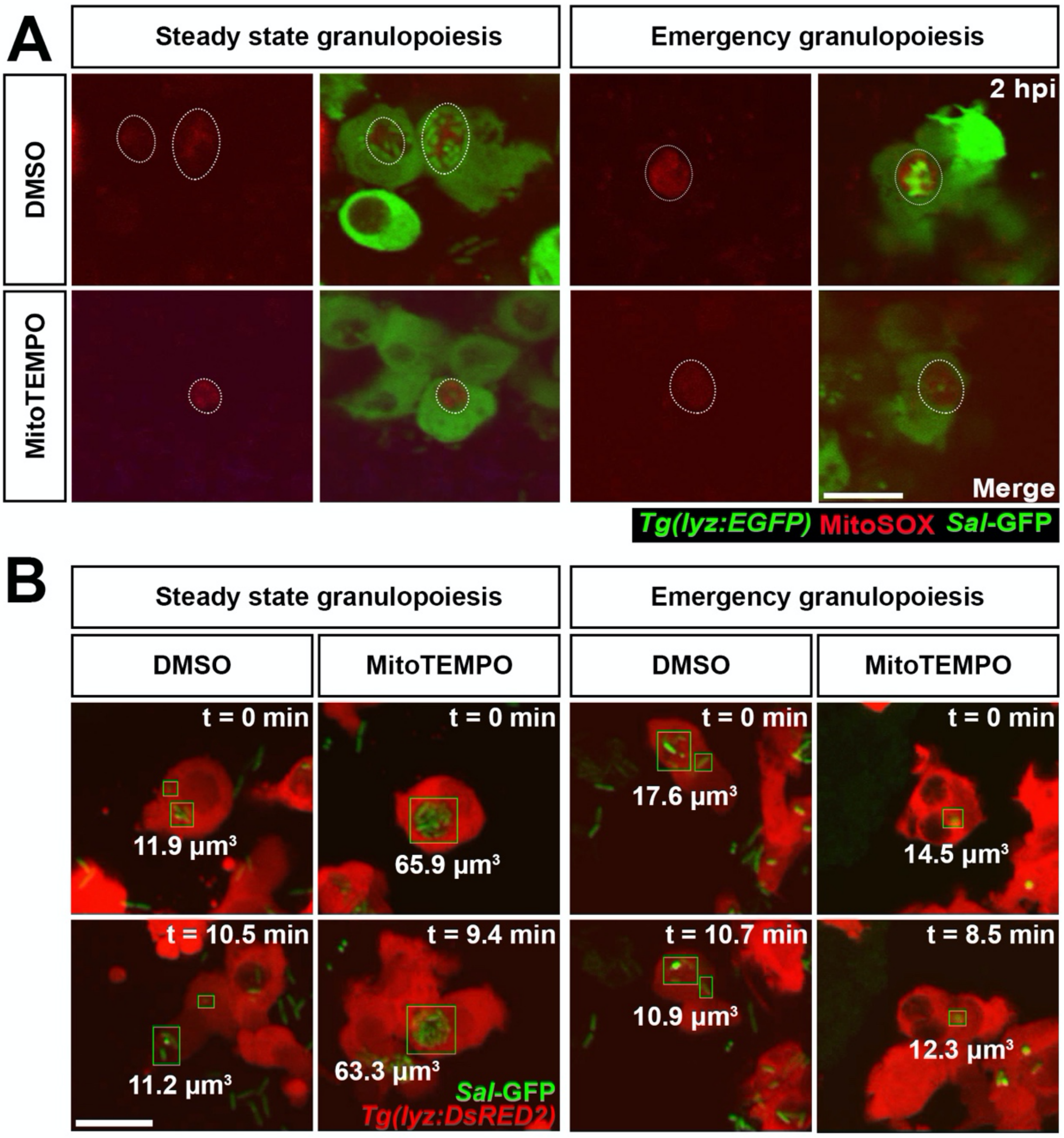


**Figure S3. Confocal imaging of mtROS production within neutrophils and killing rates following MitoTEMPO treatment. Related to Figure 3.**

(A) Live confocal imaging of mtROS production within individual *Sal*-GFP-laden SS and EG neutrophils, as detected by MitoSOX fluorescence, in the presence of DMSO (control) and MitoTEMPO. White dashed lines outline MitoSOX fluorescence within neutrophils. (B) Frame shots from live time-lapse confocal imaging of SS and EG neutrophils within *Tg(lyz:DsRED2)* larvae, in the presence of DMSO (control) and MtioTEMPO, showing volumes of intracellular *Sal*-GFP at the beginning (t=0) and end of the time-lapse experiment. Scale bar 10 µm.

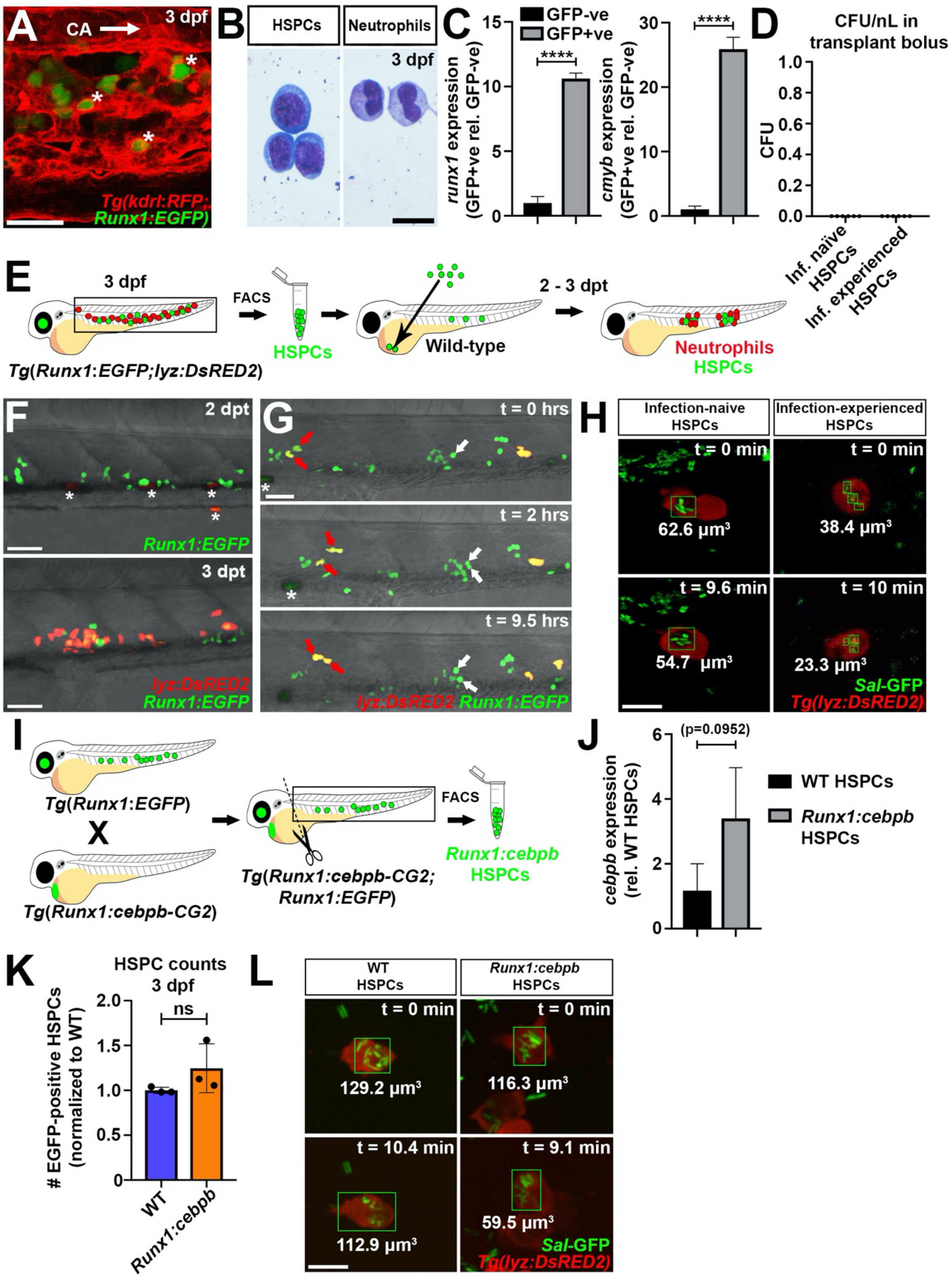


**Figure S4. The *Tg(Runx1:EGFP)* reporter line marks HSPCs, transplanted HSPCs are capable of cell division and contributing to the neutrophil lineage and confocal imaging of neutrophils derived from *cebpb*-overexpressing HSPCs. Related to Figure 4.**

(A) Live confocal imaging of HSPCs within 3 dpf *Tg(Runx1:EGFP;kdrl:RFP)* larvae showing endothelial cells ‘cuddling’ HSPCs (marked by white asterisks) in the CHT. Arrow marks the caudal artery (CA). (B) Histological examination of HSPCs and neutrophils (Wright-Giemsa stained) FACS-isolated from the trunks of 3 dpf *Tg(Runx1:EGFP;lyz:DsRED2)* larvae. (C) Expression of *runx1* and *cmyb* within GFP+ve cells (relative to GFP-ve cells) FACS-isolated from the dissected trunks of 3 dpf *Tg(Runx1:EGFP)* larvae, as detected by qPCR (in biological triplicate). (D) CFU enumeration of *Sal*-GFP within transplantation boluses (n=6 experiments). (E) Schematic illustrating HSPC transplantation protocol. EGFP-expressing HSPCs were FACS-isolated from the dissected trunks of 3 dpf *Tg(Runx1:EGFP;lyz:DsRED2)* larvae and transplanted into the circulation of 2 dpf WT recipient larvae. Transplanted larvae were visually inspected at 1 dpt for HSPC engraftment and DsRED2-expressing neutrophils (derived from transplanted HSPCs) were detectable from 2–3 days post transplant (dpt). In a typical transplantation experiment, 77.0% (SD ± 1.25, n=2 experiments) of injected recipient larvae showed engrafted HSPCs in the circulation/AGM/CHT at 1 dpt. Of HSPC-engrafted larvae, 46.1% (SD ± 0.25, n=2 experiments) possessed *DsRED2*-expressing neutrophils by 2 dpt, and 77% (SD ± 2.05, n=2 experiments) by 3 dpt. (F) Live confocal imaging of HSPC-engrafted larva at 2 and 3 dpt. (G) Frame shots from live confocal time-lapse of HSPC-engrafted larvae at 2 dpt demonstrating HSPC cell division (white arrows) and transplanted cells progressively expressing *DsRED2* (red arrows). White asterisks mark autofluorescent pigment. (H) Frame shots from live time-lapse confocal imaging of neutrophils derived from infection-naïve and -experienced HSPCs transplanted from *Tg(Runx1:EGFP;lyz:DsRED2)* larvae into WT recipients, showing volumes of intracellular *Sal*-GFP at the beginning (t=0) and end of the time-lapse experiment. (I) Schematic illustrating strategy to FACS isolate *Runx1:cebpb*-expressing HSPCs from the dissected trunks of *Tg(Runx1:cebpb-CG2;Runx1:EGFP)* larvae. (J) Expression of *cebpb* within WT and *Runx1:cebpb*-expressing HSPCs FACS-isolated from the dissected trunks of 3 dpf *Tg(Runx1:EGFP)* and *Tg(Runx1:cebpb-CG2;Runx1:EGFP)* larvae, respectively, as detected by qPCR (in biological triplicate). (K) Flow quantification of WT and *Runx1:cebpb*-expressing HSPCs from the dissected trunks of 3 dpf *Tg(Runx1:EGFP)* and *Tg(Runx1:cebpb-CG2;Runx1:EGFP)* larvae, respectively (n=25-50 larvae/sample in biological triplicate). (L) Frame shots from live time-lapse confocal imaging of neutrophils within *Tg(lyzDsRED2)* and

*Tg(Runx1:cebpb-CG2;lyz:DsRED2)* larvae, showing volumes of intracellular *Sal*-GFP at the beginning (t=0) and end of the time-lapse experiment. Error bars, mean ± SD; ns, not significant, ****p<0.0001; unpaired Student's t-test (C, J and K). CFU, colony-forming units. Scale bars, 50 μm in A, F and G, 10 μm in B, H and L.
